# Generating Synthetic Single Cell Data from Bulk RNA-seq Using a Pretrained Variational Autoencoder

**DOI:** 10.1101/2024.05.18.594837

**Authors:** Hyun Jae Cho, Eric Xie, Aidong Zhang, Stefan Bekiranov

## Abstract

Single cell RNA sequencing (scRNA-seq) is a powerful approach which generates genome-wide gene expression profiles at single cell resolution. Among its many applications, it enables determination of the transcriptional states of distinct cell types in complex tissues, thereby allowing the precise cell type and set of genes driving a disease to be identified. However, scRNA-seq remains costly, and there are extremely limited samples generated in even the most extensive human disease studies. In sharp contrast, there is a wealth of publicly available bulk RNA-seq data, in which single cell and cell type information are effectively averaged. To further leverage this wealth of RNA-seq data, methods have been developed to infer the fraction of cell types from bulk RNA-seq data using single cell data to train models. Additionally, generative AI models have been developed to generate more of an existing scRNA-seq dataset. In this study, we develop an innovative framework that takes full advantage of powerful generative AI approaches and existing scRNA-seq data to generate representative scRNA-seq data from bulk RNA-seq. Our bulk to single cell variational autoencoder-based model, termed **bulk2sc**, is trained to deconvolve pseudo-bulk RNA-seq datasets back into their constituent single-cell transcriptomes by learning the specific distributions and proportions related to each cell type. We assess the performance of bulk2sc by comparing synthetically generated scRNA-seq to actual scRNA-seq data. Application of bulk2sc to large-scale bulk RNA-seq human disease datasets could yield single cell level insights into disease processes and suggest targeted scRNA-seq experiments.

## Introduction

We are in an era where the great promise of generative artificial intelligence (AI) in synthesizing vast amounts of information and providing augmented, representative data are evident to researchers in the field and the general public. There are also many challenges in making these approaches learn and generate accurate representations of data coming from an underlying, unknown distribution. While the release of large language models including GPT-4 and Bard have made the promise and challenges (e.g., hallucination) of generative AI applied to text data clear, this is also the case for biomedical data (27) including functional genomics data. With the advent of single cell genomics, we are now capable of measuring RNA levels, open chromatin, and chromatin-chromatin interactions on a genomic scale at single cell resolution. Especially for the case of single cell RNA sequencing (scRNA-seq), this has enabled a number of applications such as identification of cell types including novel ones in complex tissues (16), interactions between cells (23) (e.g., cancer and immune cells in tumor microenvironments) and characterization of developmental trajectories within tissues harboring stem or precursor and differentiated cells (2). To get this rich set of biological insights from scRNA-seq data, there are many technical challenges to overcome including intrinsic and extrinsic technical and biological noise, undersampling of low expression and complex batch effects (4).

Taking advantage of the ability of deep learning models to filter noise, handle data sparsity and reduce data dimensionality while retaining signal in latent space, a number of powerful machine learning approaches have been developed for scRNA-seq data analysis (see (4) for a comprehensive review). Applications include improving inference of gene expression (3), identifying cell types (13), batch correction (20), and clustering (29; 10). While useful, these approaches do not address a major challenge with single cell data. It remains costly, and the number of samples in even the most extensive study remain modest. These approaches also do not take advantage of a significant opportunity. There is a wealth of bulk RNA-seq and a growing repository of scRNA-seq data. For example, as of this writing, the Trans-Omics for Precision Medicine (TOPMed) program contains *∼*69,500 RNA-seq samples. Taking advantage of this opportunity, deep learning-based, and in particular, generative AI methods could address this central challenge and make a significant contribution to the biomedical field. Specifically, development of generative AI models that generate realistic, representative scRNA-seq data from bulk RNA-seq data would enable exploration of disease states including cardiovascular disease and cancer at the single cell level for tens of thousands of patient samples as opposed to current numbers of tens to hundreds. These explorations would include downstream analysis of the generated scRNA-seq data including associating cell type abundance, cell type specific genes and pathways and developmental trajectories with disease state. Discoveries from these analyses could result in targeted single cell sequencing of specific samples for validation of the generative AI-derived results as well as further discovery and data mining.

Statistical (28; 7; 33; 32) and machine learning (26) approaches have been developed to estimate cell type proportions and average gene expression profiles from bulk RNA-seq data. Generative adversarial networks (GANs) (25), variational autoencoders (VAEs) (12) and a mixture of both (35) have been developed to augment existing scRNA-seq used as input to the networks and predict single cell gene expression response to perturbations including drug treatment. As of this writing, we recently discovered one framework, Bulk2Space (21), that generates scRNA-seq from bulk RNA-seq data and spatially maps these cells using a single cell RNA-seq and spatial reference, respectively. The scRNA-seq reference is used to train a *β*-VAE which generates the single cell data. In this paper, we develop a novel framework that generates representative scRNA-seq from bulk RNA-seq data that is a natural extension of and builds on these previous approaches. Specifically, we develop a bulk to single cell framework (termed **bulk2sc**) which utilizes a Gaussian mixture variational autoencoder (GMVAE) to generate representative, synthetic single cell data from bulk RNA-seq data by learning the cell type-specific means, variances, and proportions. **bulk2sc** is composed of three parts: a single cell GMVAE (scGMVAE) that learns cell type specific Gaussian parameters, a bulk RNA-seq VAE (Bulk VAE) that learns the cell type specific means, variances and proportion (passed from the scGMVAE) using bulk RNA-seq data as input and learned to reconstruct the scRNA data, and a bulk to single cell encoder-decoder (genVAE) composed of the encoder-decoder components from Bulk VAE that generates the synthetic, representative scRNA-seq from bulk RNA-seq data. We assess the performance of **bulk2sc** by comparing the actual, reconstructed (from scGMVAE) and synthetic scRNA-seq (from genVAE) data using qualitative (i.e., UMAP plots) and quantitative measures. Our contributions are summarized as follows:

- Once trained, our **bulk2sc** encoder-decoder is the only one that does not require further training for a given tissue type (e.g., PBMCs derived from blood) and can be applied directly using bulk RNA-seq derived from that tissue type as input to generate representative scRNA-seq data.
- The **bulk2sc** Bulk VAE is capable of learning cell type proportions, as demonstrated previously (26), as well as distributional information for each cell type including mean and variance, from pseudo-bulk RNA-seq data using supervised learning, which has never been shown.
- The success of our **bulk2sc** framework demonstrates that gene expression information can be encoded into a deep neural network encoder-decoder in such a way that using bulk RNA-seq data, in which cell type specific cell information is averaged away, as input allows faithful generation of single cell expression profiles.

## Related Work

As reviewed in (4), a number of deep learning approaches including autoencoders, variational autoencoders, Gaussian mixture variational autoencoders, graph autoencoders, adversarial autoencoders, generative adversarial networks and more have been developed for a wide variety of single cell RNA sequencing data analysis applications including imputation and denoising; removal of doublets, cell cycle variance and batch effects; dimensionality reduction; cell clustering; cell type annotation; trajectory analysis; and complete end-to-end analysis frameworks. While we recently learned of a framework, Bulk2Space (21) that generates representative scRNA-seq from bulk RNA-seq data, a number of other deep learning approaches have been developed that inform the development of our **bulk2sc** framework.

Of particular relevance to this study, a number of Gaussian mixture variational autoencoders (GMVAEs) have been developed for scRNA-seq to cluster the data according to cell types in latent space (10; 34; 22) as well as imputation, generation of cell type specific gene networks and improvement of cell developmental trajectories (34). Another deep learning approach that informs our **bulk2sc** framework is Scaden (26), in which scRNA-seq data, for which annotated cell type proportions are known, is sampled and used to generated pseudo-bulk RNA-seq as input to an ensemble neural network model which is trained to learn the cell type proportions. In this way, Scaden is trained to take bulk RNA-seq data as input and generate cell type proportions. We generalize this approach as the second step of our **bulk2sc** framework and train a deep neural network model (i.e., Bulk VAE) to learn the cell type specific means, variances and proportions, which were learned by our scGMVAE, using pseudo-bulk RNA-seq generated from the same scRNA-seq data used to train our scGMVAE. Finally, another set of relevant deep learning-based single cell approaches have been generative adversarial networks (GANs) for scRNA-seq data augmentation (12; 25; 1). These have been developed to generate more of an existing scRNA-seq dataset which is directly used as the input to train these models. These approaches and the performance metrics used to compare actual and generated/synthetic scRNA-seq data have yielded valuable insights in the development of our **bulk2sc** framework.

The Bulk2Space framework (21) is a highly relevant work as during step 1 of its training and data generation process, it generates representative scRNA-seq data from bulk RNA-seq data. During step 2, it spatially maps these single cells using a spatial reference. In step 1, a single cell reference is used to derive cell type specific means of each gene’s expression. Using the fact that bulk RNA-seq expression values (divided by the total number of cells) should be equal to the sum over cell types of cell type specific mean expression times each cell type’s proportion, the cell type proportions are derived. Further use of the same reference scRNA-seq to train a *β*-VAE allows representative scRNA-seq data to be generated according to the derived cell type proportions. This approach differs from the **bulk2sc** framework in a few important ways. Our **bulk2sc** scGMVAE explicitly learns the cell type specific distributional information including proportions, means and variances from a reference scRNA-seq dataset and trains an encoder to learn them from bulk RNA-seq data directly. This allows a stand alone bulk RNA-seq encoder to scRNA-seq decoder (genVAE) to be derived that does not require additional training for a given tissue type (e.g., PBMCs derived from blood).

## Method

Our **bulk2sc** model consists of three main components: (1) Single Cell Gaussian-Mixture Variational Autoencoder (scGMVAE), which reconstructs input scRNA-seq data by learning distributional and Gaussian mixture parameters in the latent space, (2) Bulk RNA-seq VAE which learns to decompose bulk RNA-seq data using the single cell RNA-seq-specific latent parameters learned in the previous step, and (3) Bulk to Single Cell (genVAE) VAE, which generates single cell data from bulk RNA-seq data. The schematic overview of our approach is presented in Figure 1.

**Fig. 1.**
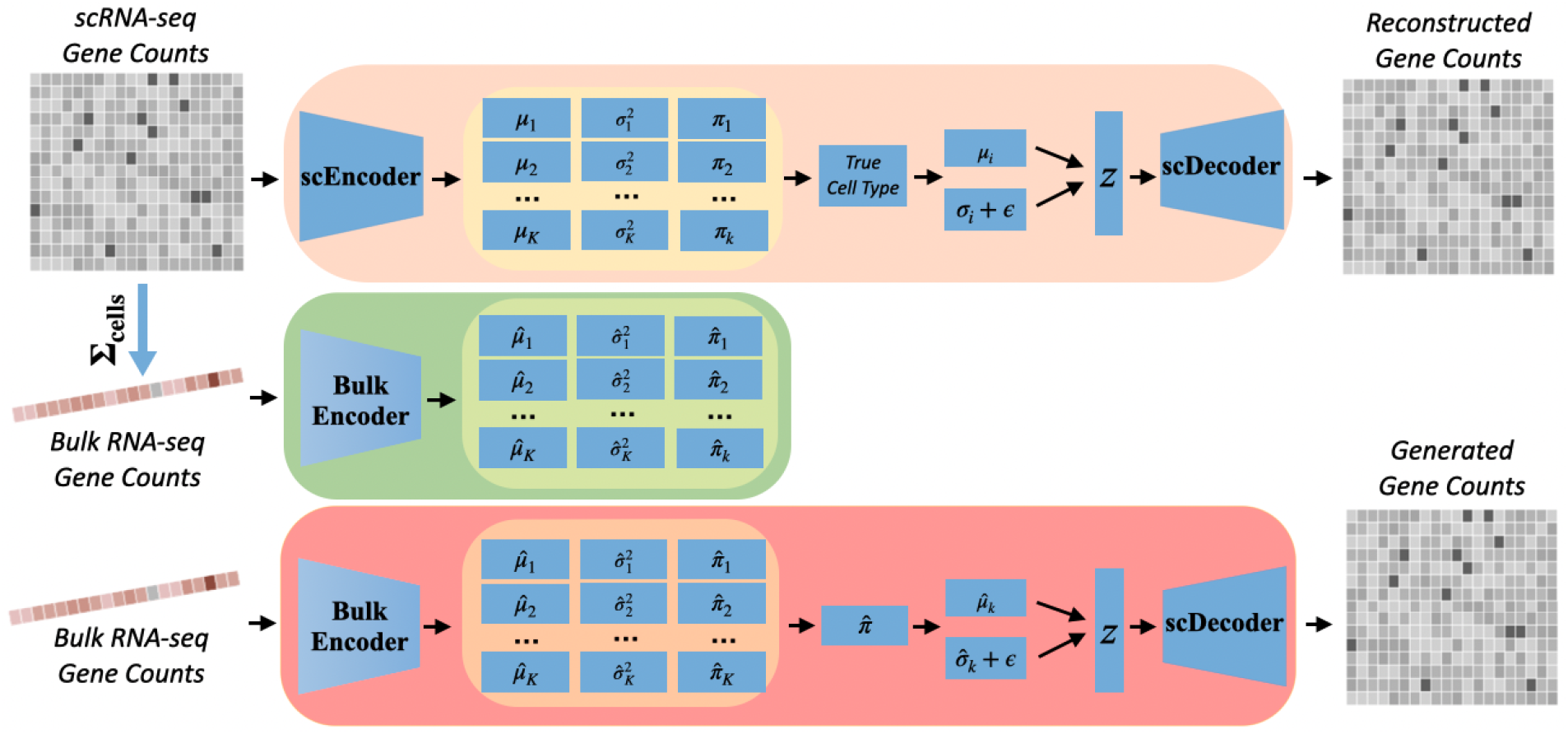
A schematic representation of the process for generating single cell data from bulk gene expression data using **bulk2sc**. Top: scGMVAE. Middle: Bulk VAE. Bottom: genVAE. Starting from single-cell RNA sequencing (scRNA-seq) gene counts, the data undergoes encoding via scEncoder, followed by Gaussian Mixture Model (GMM) processing. The data is then trained through scDecoder to produce reconstructed gene counts. Subsequently, bulk RNA-seq gene counts are processed through Bulk VAE to learn single cell-specific parameters as a function of bulk RNA-seq data. The trained model weights in Bulk VAE are transferred to genVAE to generate single cell data from bulk RNA-seq data.

### Data Preprocessing

Peripheral Blood Mononuclear Cells (PBMCs), a heterogeneous mix of blood cells all with crucial roles in the immune system, includes lymphocytes (T cells, B cells, NK cells), monocytes, and dendritic cells. Due to their abundance and accessibility, PBMCs are frequently selected for a wide range of research applications.

In this context, our method, **bulk2sc**, is specifically designed to generate realistic PBMC single-cell RNA sequencing (scRNA-seq) data by leveraging existing libraries of bulk PBMC RNA-seq data. For training **bulk2sc**, we utilized the 3K, 10K, and 68K PBMC datasets available on the 10X Genomics website. These datasets contain approximately 3,000, 10,000, and 68,000 cells, respectively, each profiling around 33,000 genes. The composition of cell types and gene sets shows slight variations across these datasets. Cell type annotations within these datasets were performed using scType (14).

To rigorously evaluate the model’s proficiency in generating single-cell data from bulk RNA-seq data, we partition the datasets into training and testing subsets, allocating 80% for training and 20% for testing. Subsequently, we employ the test set to construct pseudo-bulk data, which is then processed through genVAE. This approach enables the generation of single-cell data from unseen pseudo-bulk data.

In order to reduce remaining batch effects and to integrate multiple scRNA-seq datasets, we tried to apply Harmony (18), which effectively aligns cells across varying conditions or batches, ensuring the preservation of intrinsic biological variation. The algorithm operates on data that has been projected onto principal components (PCs), which are linearly uncorrelated variables obtained through Principal Component Analysis (PCA). However, this approach presents two following challenges for our downstream task, which requires processing bulk RNA-seq data to generate single-cell data. First, bulk RNA-seq lacks the cell-specific resolution of scRNA-seq, essential for PCA. Second, applying PCA to a single data point from bulk RNA-seq is ineffective, as PCA requires variance across multiple observations to identify meaningful components.

Additionally, it is important to note that standard data preprocessing techniques commonly applied in scRNA-seq analysis, such as normalization, scaling, and the filtering of highly variable genes and cells based on minimum gene counts, face applicability challenges in the context of single sample bulk RNA-seq data. These techniques, although crucial in contexts other than reconstruction and synthetic generation of scRNA-seq data, face transferability issues when applied to downstream single sample bulk RNA-seq data.

### Single Cell Gaussian Mixture Variational AutoEncoder: scGMVAE

The first component of our architecture, Single Cell Gaussian Mixture Variational AutoEncoder (scGMVAE), was inspired by previous works that have been developed for clustering scRNA-seq data according to cell type-specific distributions in the latent space (10; 34; 22) and is presented as the first layer in the schematic representation in Figure 1. As shown, the workflow begins with the scRNA-seq gene counts. scEncoder takes in the input count data and captures essential latent features, quantifying them into two main components: cell type (*k*)-specific mean (*μ*_*k*_) and variance 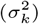. In addition, scEncoder learns the cell type proportion (*π*_*k*_). We encoded every PBMC cell type in scType, with a total of 34 distinct cell types (*K*=34.) With the inclusion of Gaussian noise *ϵ*_*i*_, the latent representation *z* is then reparameterized as follows: 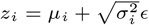, where *i* represents the index of the input cell. *z* is then used as the input to the scDecoder model, which tries to reconstruct the input scRNA-seq count data by leveraging two types of reconstruction loss functions: Mean Squared Error (MSE) and Zero-Inflated Negative Binomial (ZINB). For the input count data *X* and the reconstructed data 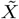, the MSE is computed:

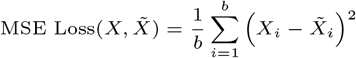

for batch size *b*. Finally, for computing the ZINB loss, scDecoder outputs two parameters in addition to the reconstruction: the over-dispersion parameter *r* and the zero-inflation parameter *θ*, and the ZINB loss is computed in the following way:

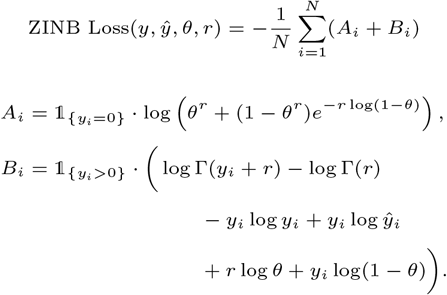

Finally, we adjust the typical Kullback–Leibler (KL) divergence (19) to account for the mixture of *K* Gaussian prior distributions as follows:

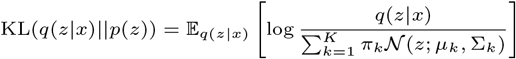

where *q*(*z*|*x*) is the posterior distribution of the latent variable *z* given the input *x ∼ X*; *p*(*z*) is the GMM prior; *π*_*k*_ represents the mixing coefficients for the Gaussian components (learned cell type proportion), summing up to 1; 𝒩 (*z*; *μ*_*k*_, Σ_*k*_) is the *k*-th Gaussian component of the mixture, with mean *μ*_*k*_ and covariance Σ_*k*_; 𝔼_*q*(*z*|*x*)_[*·*] denotes the expectation with respect to the posterior distribution. To prevent posterior collapse, we clip the value of KL divergence. The aggregate loss function for scGMVAE becomes the sum of the three loss terms, as follows.

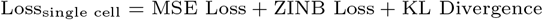

### Bulk RNA-seq Encoder: Bulk Encoder

Bulk Encoder is the intermediate layer in Figure 1, and it is the critical component that learns the cell type-specific estimates of their proportions, means, variance 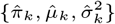 as a function of bulk RNA-seq data. Bulk RNA-seq data is created by sampling cells from the scRNA-seq data and summing the expression values to create pseudo-bulk expression data. Importantly, we hypothesize that while bulk or pseudo-bulk RNA-seq expression values for each gene, *x*^*b*^, represent the following weighted average over cell types, 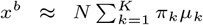 (where *N* is the total number of single cells), there are genes and gene expression profiles that are specific to cell types and contain enough information to enable cell type-specific distributional parameters to be learned from bulk RNA-seq data. We were able to verify this by a rapid decrease in the MSE between the GMM parameters and those learned by Bulk Encoder. The learning of the parameters is accomplished by taking the MSE between the output parameters 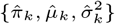 and the corresponding parameters {*π*_*k*_, *μ*_*k*_, 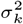} from the scGMVAE model:

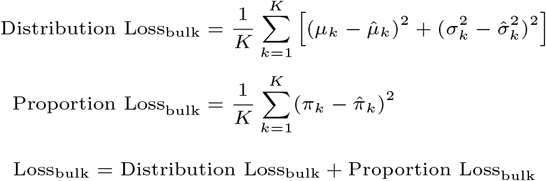

### Bulk to Single Cell Generator: genVAE

The final component of our model architecture is called genVAE, and it is a VAE model that is responsible for generating single cell data from bulk RNA-seq input data, as shown in the lowermost layer in Figure 1. As the figure indicates, the trained parameters from Bulk Encoder and scDecoder are transferred to the encoder and the decoder of genVAE respectively.

The process of generating single cells utilizes bulk RNA-seq data as an input for the Bulk Encoder, which produces cell type-specific parameters: 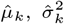, and 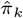. To ensure that the generated cells reflect the proportions of cell types present in the input data, we employ 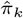 as a weighted factor to sample 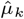, and 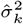. Consequently, we reparameterize to get a latent variable *z*:

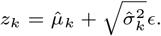

Each *z*_*k*_ is used to generate one single cell that resembles cells with type *k*; therefore, reparameterizing *m* times will generate *m* cells that follow the distribution learned in 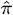.

For detailed model parameter settings, please refer to our Github link, provided in the Data and Source Code Availability section.

### Qualitative Evaluation Metrics

We categorize our evaluation of the **bulk2sc** model into qualitative and quantitative metrics to comprehensively measure its performance. Specifically, for qualitatively assessing the model, we employ visualizations through Uniform Manifold Approximation and Projection (UMAP) plots at five distinct stages of our model: 1) the input count data, 2) the reparameterized latent representations, sampled from the learned parameters of the Gaussian Mixture Model (GMM), 3) the reconstructed data, 4) the data generated by genVAE from pseudo-bulk data synthesized using the training set, and 5) the data generated by genVAE from pseudo-bulk data synthesized using the testing dataset. UMAP visualizations using at different stages enable the assessment of various components of the model. Clear cell type-specific clusters in UMAPs imply that cells with similar gene expression profiles group closely together, forming distinguishable clusters. In addition, comparing cell type proportions among the real, input datasets and generated dataset provide significant insights into the model’s ability to capture the cell type-specific information from bulk RNA-seq datasets.

The second approach to our qualitative evaluations involves creating correlation heatmaps for the genes exhibiting the highest variability. This process entails selecting the top 100 genes with the greatest variation in the input count data. We then analyze the correlations between these genes, both within the input count data and in the data generated by our model. A high degree of similarity suggests that the generated data accurately reflects the underlying biological variability present in the real data.

Lastly, we employ QQ (Quantile-Quantile) plots to quantitatively evaluate the alignment of distributions across different datasets: the input data, the single cell data generated from the training data, and the single-cell data generated from the test data. These QQ plots provide a nuanced perspective on the comparative distribution between the model-generated single-cell data and the original input data which are used assess how closely the model replicates the statistical characteristics of the sparse count nature of scRNA-seq data.

### Quantitative Evaluation Metrics

Additionally, we employ a range of quantitative metrics to assess the similarity between the input data and the generated single cell data. These metrics include cosine similarity, Pearson correlation coefficient, iLISI (integration Local Inverse Simpson’s Index) (18), and a variant of Correlation Discrepancy (12). Cosine similarity focuses on the orientation of gene expression vectors, thus comparing the profiles based on patterns rather than absolute values, which is particularly useful given the variability in gene expression scales across single cells. On the other hand, Pearson’s correlation coefficient is effective in capturing co-expression patterns, thus reflecting the biological co-regulation of genes. Both cosine similarity and Pearson’s correlation coefficient are widely used to compare the overall structure in gene expression data, making them suitable for assessing the quality of synthetic scRNA-seq datasets against real ones (6; 15).

Additionally, we employ iLISI as a method to quantitatively evaluate the degree of similarity between the generated data and the original dataset. The computation of iLISI is a multi-step process, beginning with the identification of cell neighborhoods within a dimension-reduced space, achieved through PCA (Principal Component Analysis) in this analysis. This is then followed by determining the proportion of cells from each source within each neighborhood, and finally, computing the Inverse Simpson’s Index based on these proportions.

To ascertain the efficacy of our data generation method using iLISI, we apply PCA to both the generated and original datasets, then compute the iLISI value. The level of integration indicated by the score serves as a measure of the accuracy of our synthetic data in replicating the structure of the original data. It is important to note, however, that iLISI is sensitive to the size of the datasets under comparison since it relies on proportional data (18). Therefore, the maximum achievable iLISI value based on the disparity between the sizes of the datasets can be mathematically deduced as follows:

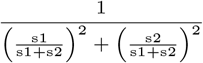

where s1 and s2 represent the sizes of the datasets used in the iLISI calculation.

In (12), an evaluation method based on the Correlation Discrepancy (CD) metric was introduced to assess the 1-norm of the difference in correlations to measure the discrepancy between the correlations in real and synthetic data, defined as:

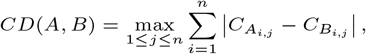

where *C*_*A*_ and *C*_*B*_ are the Spearman’s rank correlation matrices of synthetic and real data, respectively. A CD value of 0 indicates no discrepancy, with lower values denoting higher quality of the synthetic data in preserving gene-gene correlations. The worst value of CD occurs when the differences between the correlation coefficients in matrices A and B are at their maximum possible value. Since correlation coefficients range between -1 and 1, the maximum absolute difference between two correlation coefficients is 2. Given an *n × n* correlation matrix with *n* indicating the *n* most highly variable genes, a value of CD(AB) = 2(*n −* 1) would indicate the maximum discrepancy, where *n −* 1 accounts for the diagonal. In our study, we apply the CD metric to the 100 most highly variable genes.

While the Pearson correlation heatmaps offer a qualitative insight into the relationships between genes, the CD metric provides a more nuanced and quantitative assessment of the similarity between the synthetic and real data. This dual approach allows for a comprehensive evaluation of the fidelity of the synthetic data in mirroring the complex gene-gene interaction patterns found in the original scRNA-seq data.

## Experimental Results

### Qualitative Results

For each PBMC dataset, we created five UMAPs representing different stages: the real input scRNA-seq data, the reparameterized *z* values derived from *μ*_*k*_ and 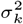, scGMVAE reconstructed data, and genVAE processed cells from train and test set pseudo-bulk data.

Figures 2, 4 (in Supplementary Materials), and 5 (in Supplementary Materials) showcase UMAP visualizations corresponding to the five distinct stages of our analysis for the 3K, 10K, and 68K PBMC datasets, respectively. Panel a of each figure displays the input scRNA-seq data with color-coded cell types. These datasets show distinct yet partially overlapping cell type clusters and certain cell types are characterized by more than one distinct cluster.

**Fig. 2.**
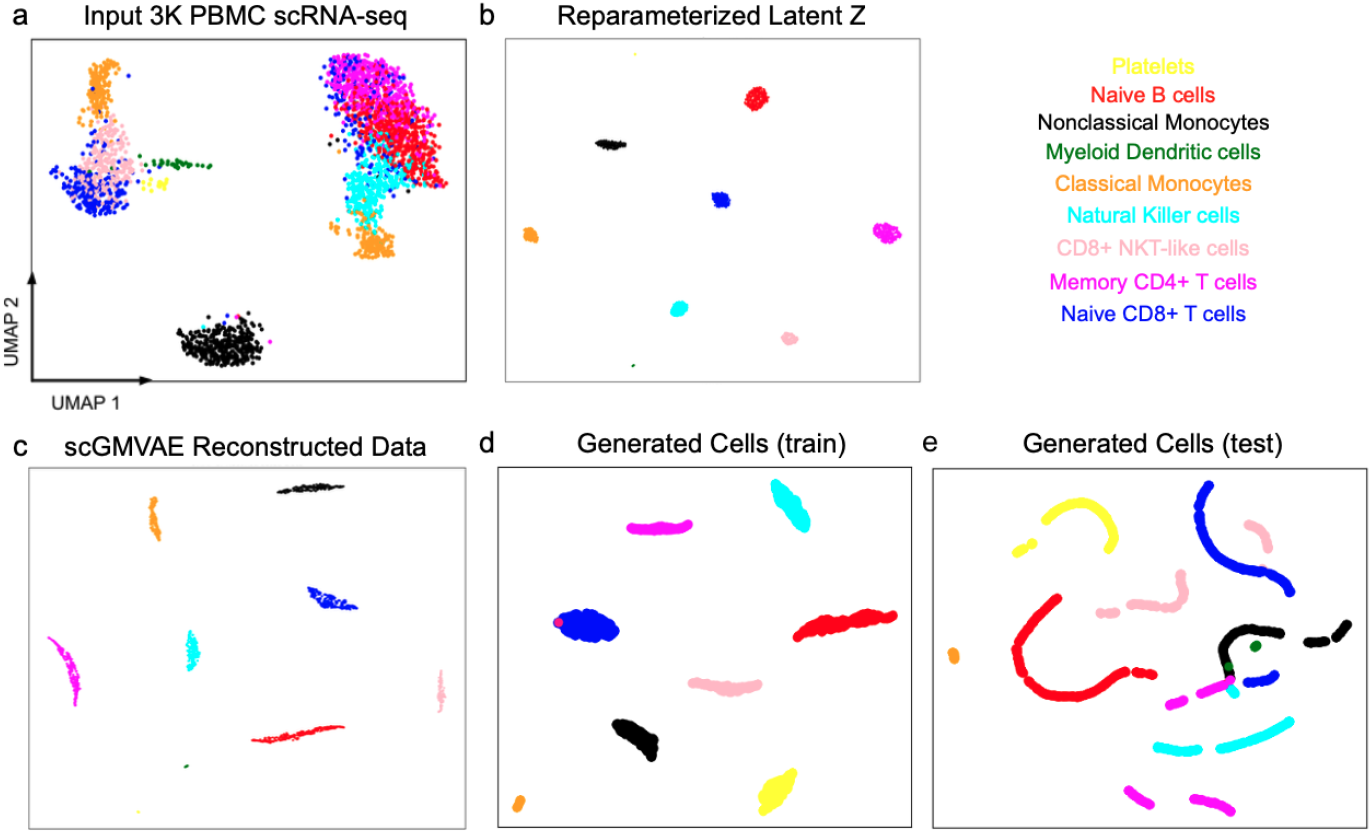
3K PBMC visualizations through UMAP plots at five distinct stages of our model. a. the input count data, b. the reparameterized latent representations, sampled from the learned parameters of the Gaussian Mixture Model (GMM), c. the reconstructed data, d. the data generated by genVAE from pseudo-bulk data synthesized using the training set, and e. the data generated by genVAE from pseudo-bulk data synthesized using the testing dataset.

**Fig. 3.**
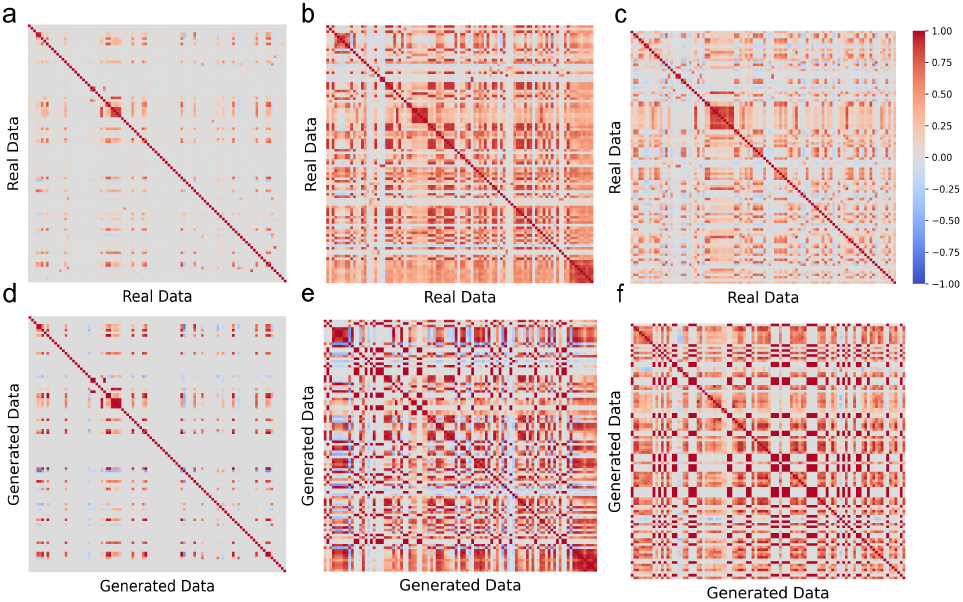
Visual representation of Pearson correlations among the 100 most variable genes. a-c: Real PBMC data – 3K, 10K, and 68K cells respectively. d-f: generated data – 3K, 10K, and 68K cells respectively. These heatmaps illustrate the similarity of gene-to-gene relations that are captured by the generated cells. Similar maps imply better matching of the gene-to-gene relations between the real and generated cells.

**Fig. 4.**
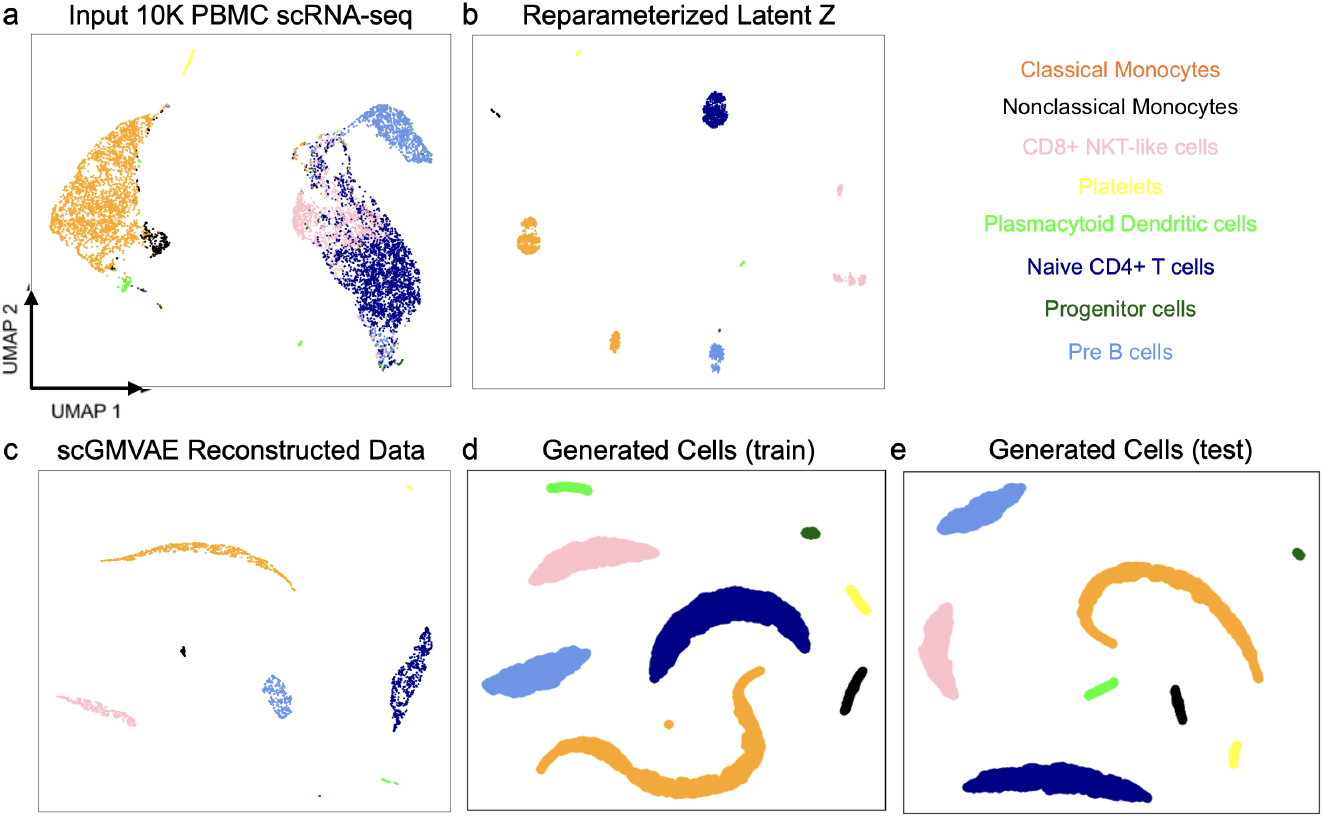
10K PBMC visualizations through UMAP plots at five distinct stages of our model. a. the input count data, b. the reparameterized latent representations, sampled from the learned parameters of the Gaussian Mixture Model (GMM), c. the reconstructed data, d. the data generated by genVAE from pseudo-bulk data synthesized using the training set, and e. the data generated by genVAE from pseudo-bulk data synthesized using the testing dataset.

**Fig. 5.**
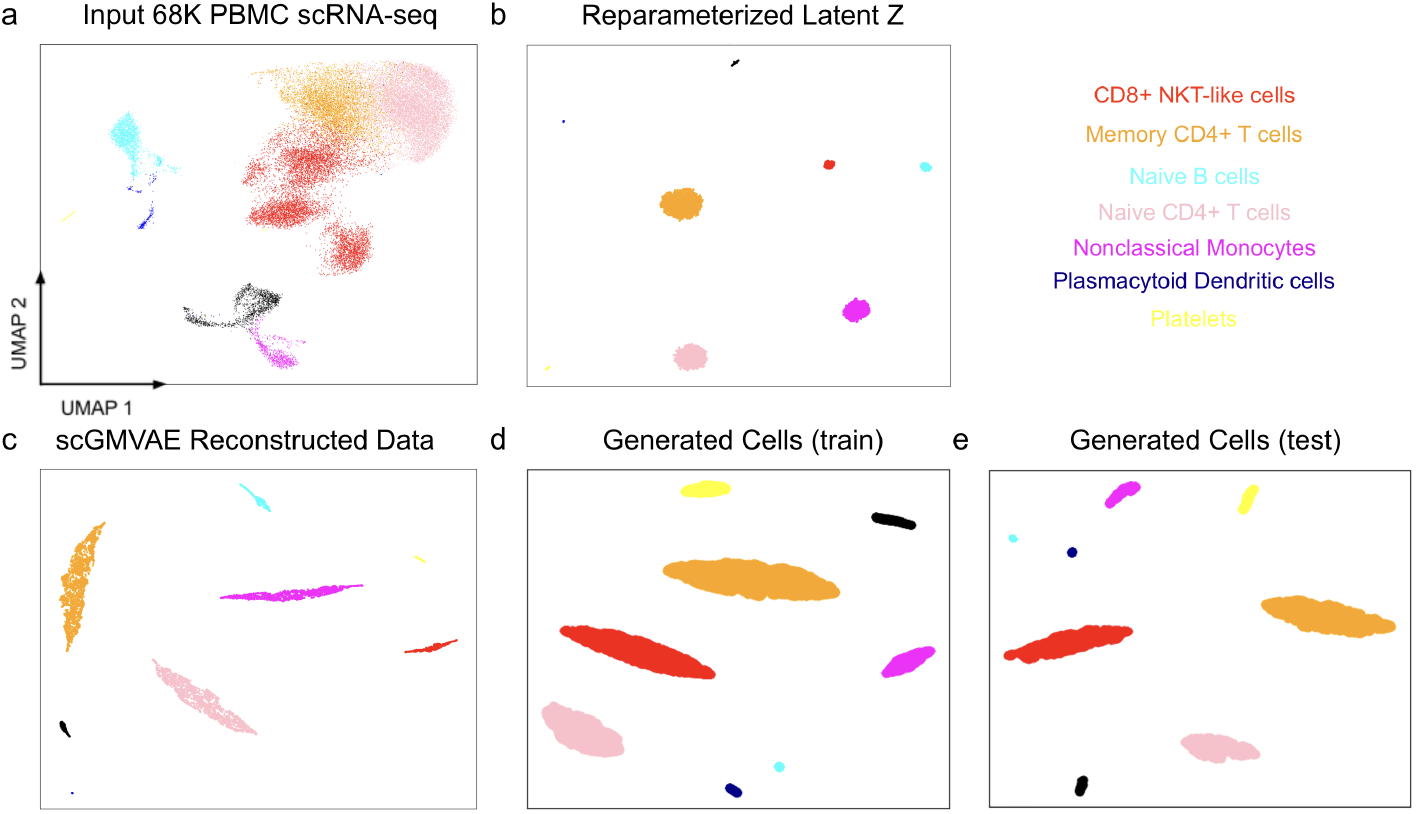
68K PBMC visualizations through UMAP plots at five distinct stages of our model. a. the input count data, b. the reparameterized latent representations, sampled from the learned parameters of the Gaussian Mixture Model (GMM), c. the reconstructed data, d. the data generated by genVAE from pseudo-bulk data synthesized using the training set, and e. the data generated by genVAE from pseudo-bulk data synthesized using the testing dataset.

Each panel b displays reparameterized latent variable *z*, notably arranged in a manner suggestive of Gaussian distribution patterns, affirming our model’s assumption of Gaussian distributions in the latent space. This indicates that the reparameterization effectively captures the probabilistic nature of the data.

Each panel c shows the reconstructed data from scDecoder. Because scGMVAE model learned cell type-specific distribution parameters *μ*_*k*_ and *Σ*_*k*_, the cell clusters are more unique than in the original dataset. In both panels b and c, we observe that the cell clusters contain data points with approximately similar proportions.

Finally, panels d and e visualize generated cells derived from the pseudo-bulk of the training and testing sets, respectively. These plots reveal distinct cell type-specific clusters similar to original scRNA-seq data. In addition, as observed in panels b and c, the proportions of these clusters in the generated data closely align with those in the original dataset, and the comparisons between the cell type-proportions in the scRNA-seq data and the the generated single cell data is highlighted in Figure 6 in Supplementary Materials. This resemblance is indicative of the model’s ability to not only generate cells with the type-specific characteristics accurately but also to maintain the relative abundance of each cell type as found in the original data.

**Fig. 6.**
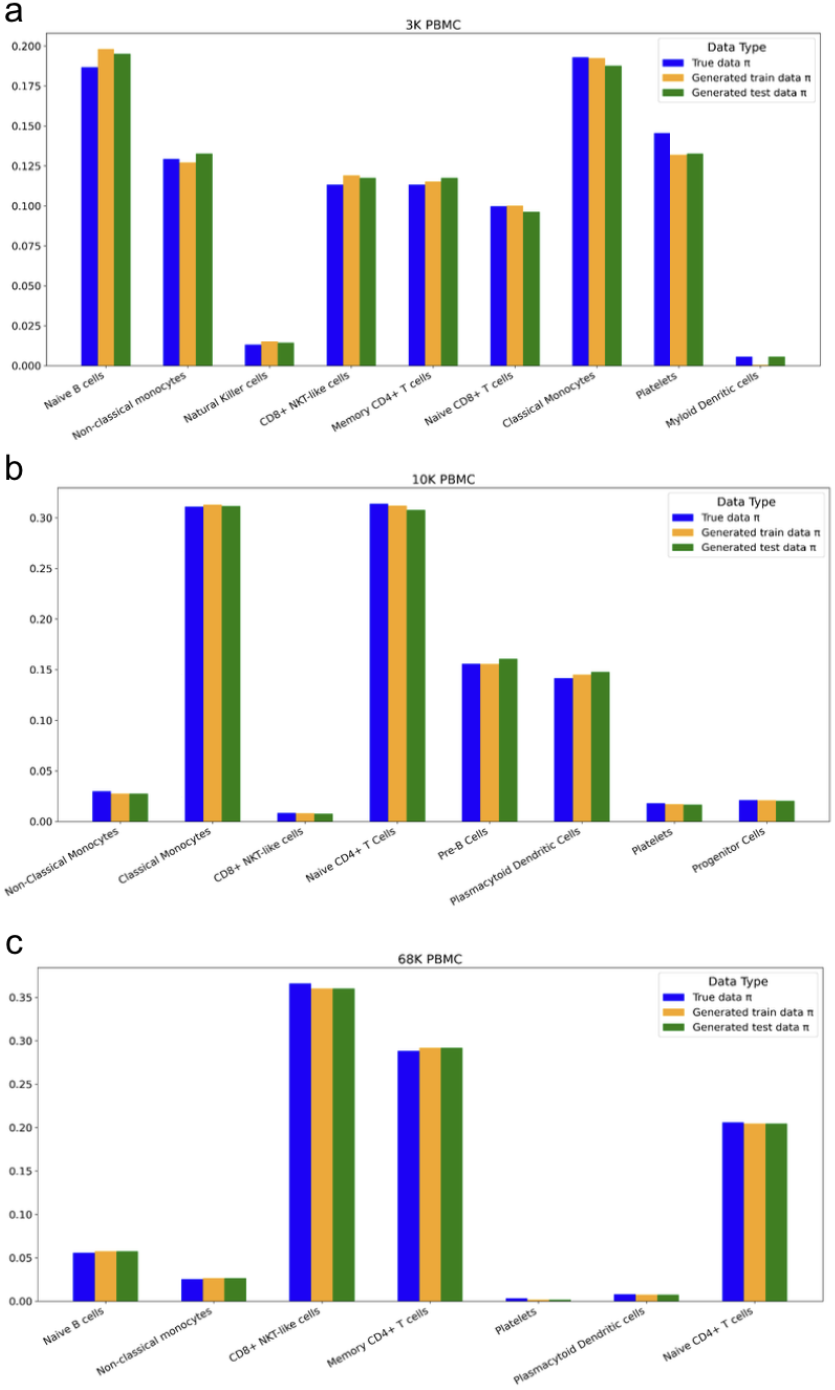
Cell type proportions of input data (blue), generated data from training data (orange), generated data from testing data (green). a. 3K PBMC. b. 10K PBMC. c. 68K PBMC. x-axis represents cell types and y-axis represents proportions.

Figure 3 displays the heatmaps representing Pearson correlations among the 100 most variable genes. The gene selection process involved isolating the top 10% genes with the highest mean expression, followed by identifying the 100 genes exhibiting the highest variance-to-mean ratio from this subset in the input scRNA-seq dataset in order to capture genes with both elevated average expression levels and substantial variability across the samples. The heatmaps derived from real PBMC data (panels a-c) compared to the heatmaps derived from the generated PBMC data (panels d-f) show notable similarities in the overall trends and patterns of gene-to-gene relationships. It is also noticable that the heatmaps of the generated data seem to exhibit correlations with more pronounced and intensified values overall, which could indicate a slight exaggeration of the correlations between genes among the generated cells.

In Supplementary Materials, Figure 7 showcases Quantile-Quantile (QQ) plots that provide insights into the distributional characteristics of both scRNA-seq and generated data. Specifically, panels a, d, and g display the average gene counts for the 3K, 10K, and 68K PBMC scRNA-seq datasets, respectively. Panels b, e, and h show the average gene counts for scRNA-seq data generated from pseudo-bulk RNA-seq data using the training set. Meanwhile, panels c, f, and i illustrate the average gene counts for data generated using the test set. The remarkable similarities in the distributional characteristics across these three sets of plots highlight the model’s proficiency in generating single-cell data that closely mirrors the gene counts of the actual input data.

**Fig. 7.**
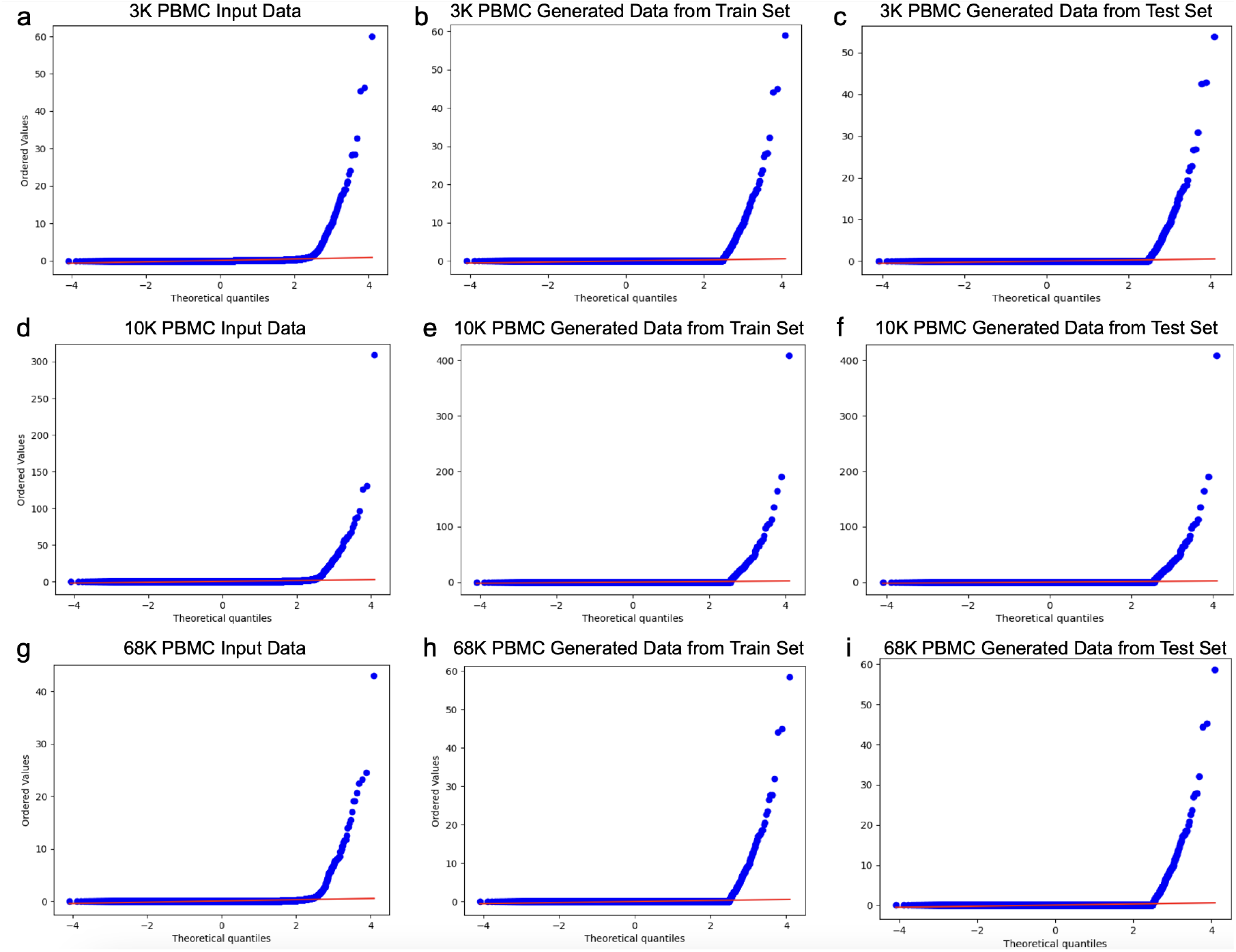
Quantile-Quantile (QQ) plots demonstrating the distributional characteristics of scRNA-seq data across different datasets and conditions. Panels a, d, and g illustrate the QQ plots for the original scRNA-seq mean count values at 3K, 10K, and 68K PBMC input sizes, respectively. Panels b, e, and h display the QQ plots for the mean count values of scRNA-seq data generated from pseudo-bulk RNA-seq data using the training set, corresponding to the same 3K, 10K, and 68K PBMC sizes. Panels c, f, and i show the QQ plots for the mean count values in scRNA-seq data generated from pseudo-bulk RNA-seq data using the test set, again matched to the 3K, 10K, and 68K PBMC sizes. These plots are crucial for comparing the distributional alignment between the original scRNA-seq data and the model-generated data from both the training and test sets, providing insights into the model’s ability to replicate distributional characteristics of the original scRNA-seq data.

In summary, the qualitative assessment methods we’ve detailed, including UMAPs (Uniform Manifold Approximation and Projection), heatmap comparisons, and QQ (Quantile-Quantile) plots, together offer a detailed view of the relationship between single cells generated from pseudo-bulk datasets and actual scRNA-seq datasets. These techniques facilitate an in-depth analysis, emphasizing the capability of the **bulk2sc** model in accurately mirroring the features of real single-cell RNA sequencing data based on bulk data inputs.

### Quantitative Results

In Table 1, we provide the cosine similarity scores among the input scRNA-seq, the reconstructed and the generated data. Remarkably, each of these scores registers at or exceeds the threshold of 0.96. While the inherent sparsity of the datasets plays a crucial role in the cosine similarity metrics, the observed high levels of similarity provide strong evidence that the reconstructed and generated data by **bulk2sc** are remarkably congruent with the original scRNA-seq data.

**Table 1.**
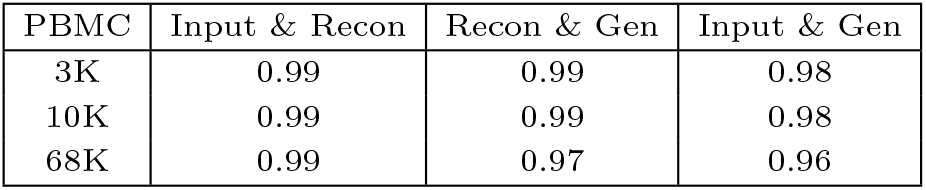
Comparisons between the input scRNA-seq data, reconstructed data, and generated cells using the average cosine similarity [-1, 1].

In Table 2, we present the average Pearson correlation coefficients between the scRNA-seq data, the data reconstructed through scGMVAE, and the cells generated using genVAE. Although the results are varied between values of 0.77 and 0.99, the computed correlations show relatively strong positive correlations that underscore a robust and consistent linear relationship between each pair of datasets.

**Table 2.** Comparisons between the input scRNA-seq data, reconstructed data, and generated cells using the average Pearson correlation coefficient (Pearson) [-1, 1].

| PBMC | Input & Recon | Recon & Gen | Input & Gen |
| --- | --- | --- | --- |
| 3K | 0.81 | 0.91 | 0.81 |
| 10K | 0.82 | 0.77 | 0.82 |
| 68K | 0.83 | 0.99 | 0.83 |

The cosine similarities and Pearson correlations align with the QQ plots presented in Figure 7 in Supplementary Materials, which shows how much the the distributional characteristics align among the three datasets for all 3K, 10K, and 68K PBMC datasets we tested.

Finally, in Table 3, we present the results for the correlation discrepancy (CD). Contrasting the robust outcomes observed in Table 2, the CD metric reveals a more varied set of results. This variation can be largely attributed to the nature of CD’s underlying metric, Spearman’s rank correlation. Unlike Pearson’s correlation, which assesses linear relationships, Spearman’s rank correlation evaluates monotonic relationships, making it sensitive to both linear and non-linear associations. This sensitivity can be particularly relevant in the context of sparse scRNA-seq data, where gene expression patterns are often complex and do not necessarily follow linear trends.

**Table 3.**
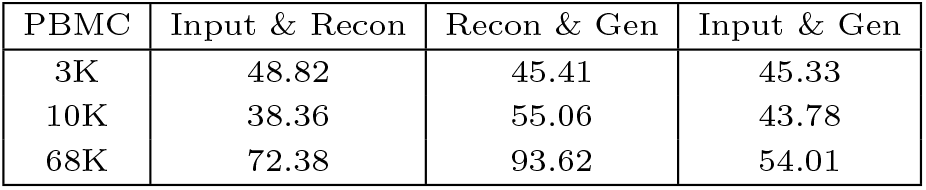
Correlation Discrepancy (CD) for comparing the gene-gene correlations between two datasets. Evaluated on 100 most highly variable genes, the range of CD is [0, 198], where 0 indicates no discrepancy (best) and 198 indicates maximum discrepancy (worst).

In Table 4, we present the Integration Local Inverse Simpson’s Index (iLISI) scores for each pairing within each dataset. Notably, the iLISI scores for training versus test dataset comparisons closely align with their theoretical capacity, indicating highly effective integration, in contrast to the lower scores observed in other dataset pairings. Furthermore, these training and test comparisons required significantly fewer principal components to achieve at least 90% variance, underscoring a simpler underlying structure and higher similarity compared to comparisons with the original dataset.

**Table 4.** Integration Local Inverse Simpson’s Index (iLISI) for Assessing Dataset Integration Quality. ‘iLISI’ represents the level of integration. ‘Capacity’ denotes the iLISI score assuming maximal integration, determined by the dataset sizes. ‘% Var’ shows the percentage of total variance explained by the selected principal components, with a minimum of 90% required for each comparison. ‘# PCs’ lists the number of principal components used.

|  | iLISI | Capacity | % Var | # PCs |
| --- | --- | --- | --- | --- |
| 3k Train & Original | 1.048 | 1.969 | 90.00 | 1638 |
| 3k Train & Test | 1.114 | 1.368 | 92.62 | 4 |
| 3k Test & Original | 1.065 | 1.290 | 90.00 | 1724 |
| 10k Train & Original | 1.016 | 1.883 | 90.00 | 3667 |
| 10k Train & Test | 1.642 | 1.647 | 93.52 | 3 |
| 10k Test & Original | 1.035 | 1.890 | 90.00 | 3993 |
| 68k Train & Original | 1.019 | 1.541 | 90.00 | 3993 |
| 68k Train & Test | 1.471 | 1.471 | 92.95 | 3 |
| 68k Test & Original | 1.023 | 1.146 | 90.00 | 4761 |

Overall, the quantitative evaluations conducted in our study establish a solid foundation for assessing the accuracy and reliability of the **bulk2sc** model. By employing a variety of statistical measures and comparison methods, these evaluations provide a comprehensive and unbiased examination of the model’s ability to replicate single-cell RNA sequencing data from pseudo-bulk data. The uniformity of findings across different metrics underscores the model’s proficiency in accurately capturing key characteristics of scRNA-seq data. This demonstrates the model’s potential as an invaluable resource for utilizing existing bulk RNA-seq datasets to generate realistic synthetic single-cell RNA-seq data, which provides more precise understanding of cellular heterogeneity, gene expression dynamics, and biological processes at the individual cell level over the bulk counterpart.

### Downstream Analysis

The primary application of **bulk2sc** lies in leveraging the abundant repository of bulk RNA-seq datasets to simulate scRNA-seq data. To assess the effectiveness of this approach, we sought a study that incorporated both bulk RNA-seq and corresponding scRNA-seq PBMC data, which was available in (8). The evaluation procedure involves initially training **bulk2sc** on the scRNA-seq data, subsequently producing single-cell data from the bulk RNA-seq data. The final step involves conducting comparative evaluations between these paired datasets, specifically contrasting the derived single-cell data with the original bulk RNA-seq data.

In the study conducted in (8), scRNA-seq methods were evaluated using PBMCs in two experimental replicates, PBMC1 and PBMC2. These replicates are essential for assessing the reproducibility of various scRNA-seq techniques. We were unable to apply scType for cell type annotation; hence, we used Leiden (31) for cell type subgroup clustering method.

In Figure 8 in Supplementary Materials, we provide UMAP visualizations of the scRNA-seq data (panel a), variation stages in training our **bulk2sc** model, such as the reparameterized latent representation *z* (panel b), the reconstructed data (panel c), and the generated single cell data from genVAE (panel d). Panels e and f represent UMAPs of single cell data generated from bulk RNA-seq data, PBMC1 and PBMC2 respectively. It is evident that the single cell data **bulk2sc** generated using both pseudo-bulk and real bulk RNA-seq datasets form clear cell type-specific clusters, suggesting that the model is capable of modeling cell type distributions as a function of bulk RNA-seq data.

**Fig. 8.**
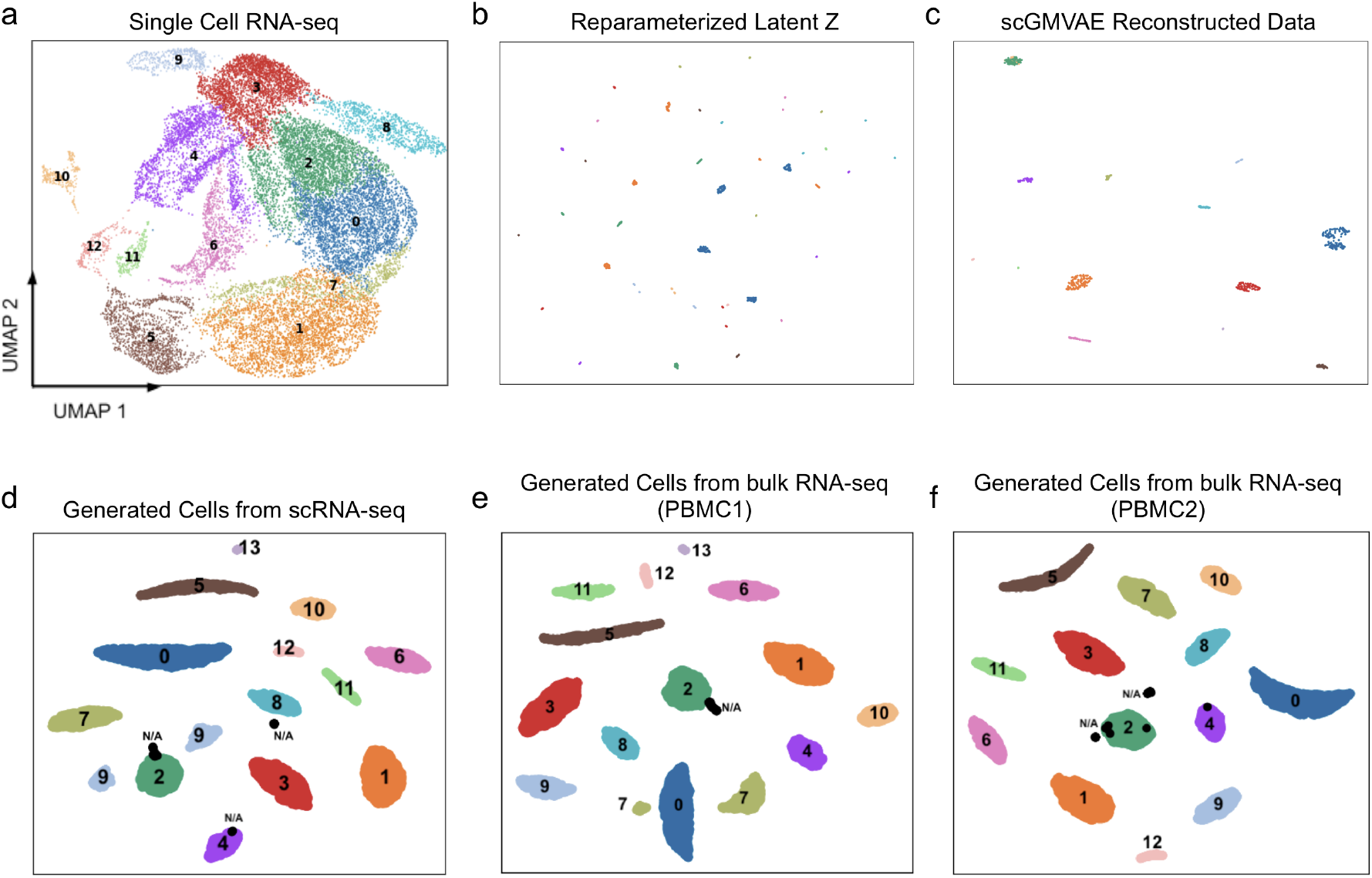
In this study, we evaluated the similarity between cells generated from bulk RNA-seq datasets and real single-cell data, using dataset pairs from Ding et al. (2020) (8). Our analysis included: a. UMAP visualization of single-cell RNA-seq data. b. UMAP of reparameterized *z*. c. UMAP of data reconstructed from the scGMVAE model. d. UMAP of cells generated from pseudo-bulk data synthesized from single-cell RNA-seq data. e. UMAP of single-cell data generated from PBMC1 bulk RNA-seq data. f. UMAP of single-cell data generated from PBMC2 bulk RNA-seq data. N/A represents cell types that were not present in the scRNA-seq dataset. Cell type subgroup annotation was performed using the Leiden clustering algorithm.

**bulk2sc**’s ability to accurately model cell type proportion is further demonstrated in Figure 9 in Supplementary Materials, which compares the cell type proportions in the scRNA-seq data (blue), the generated cells from scRNA-seq data (orange), the generated cells from PBMC1 bulk RNA-seq data (green), and the generated cells from PBMC2 bulk RNA-seq data (pink). As shown clearly, the proportions in all generated single cell data follow the proportions in the scRNA-seq data.

**Fig. 9.**
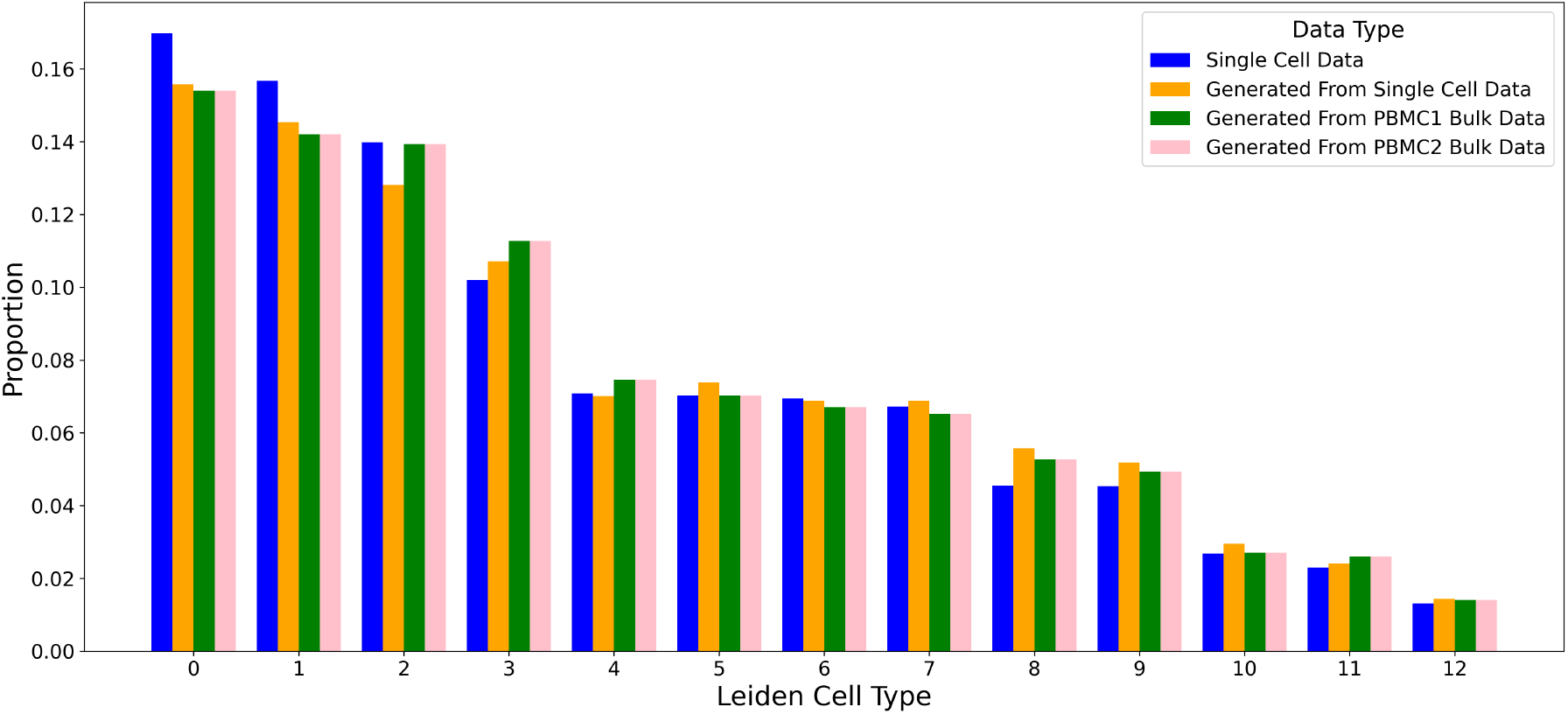
The proportions of cell types in scRNA-seq data (blue), in cells generated from pseudo-bulk of scRNA-seq (orange), in cells generated from PBMC1 bulk data (green), and in cells generated from PBMC2 bulk data (pink). The cell type proportion information is well maintained in the single cell data generated from both pseudo-bulk and real bulk PBMC RNA-seq datasets.

In Table 5 in Supplementary Materials, we delineate the cosine similarity, Pearson correlation coefficient, iLISI, and the Correlation Discrepancy between the four single cell datasets: the scRNA-seq data, the generated single cell data from the pseudo-bulk dataset created by summing the scRNA-seq data, and two generated single cell data from PBMC1 and PBMC2 true bulk RNA-seq datasets.

**Table 5.** Comparisons between the input scRNA-seq data, reconstructed data, and generated cells. scRNA-seq represents the single cell data from (8). scGenerated represents the generated cells from scRNA-seq data using trained **bulk2sc**. scPBMC1 and scPBMC2 represent single cell generated from PBMC1 bulk RNA-seq data and PBMC2 bulk RNA-seq data respectively, which are also provided by (8) as the paired bulk datasets to the scRNA-seq. Cosine represents cosine similarity, which measures the cosine of the angle between two non-zero vectors ranging in [-1, 1]. Pearson represents Pearson correlation coefficient, which assesses the linear relationship between two datasets to show the strength and direction of the association between ranges of [-1, 1]. iLISI (Integration Local Inverse Simpson’s Index), introduced in (18), measures the integration quality of combined datasets, with higher scores indicating better mixing of cells. In the table, iLISI scores are paired with their capacity, the maximum iLISI score given complete integration dictated by dataset size disparity, in the form (“score | capacity”). CD represents a derivation of Correlation Discrepancy introduced in (12), and it is computed as the 1-norm of the difference in correlations among a set of genes. We compute the CD among 100 most highly variable genes, which makes the range of CD is [0, 198], where 0 indicates no discrepancy (best) and 198 indicates maximum discrepancy (worst).

| Metric | scRNA-seq & scGenerated | scRNA-seq & scPBMC1 | scGenerated & scPBMC1 | scPBMC1 & scPBMC2 |
| --- | --- | --- | --- | --- |
| Cosine | 0.97 | 0.97 | 0.99 | 0.99 |
| Pearson | 0.98 | 0.99 | 0.99 | 0.99 |
| iLISI | 1.106 \| 1.356 | 1.110 \| 1.356 | 1.867 \| 1.984 | 1.933 \| 1.984 |
| CD | 37.72 | 37.44 | 2.34 | 3.29 |

The analysis revealed uniformly high cosine similarity scores across all dataset comparisons, with values ranging from 0.97 to 0.99. This indicates a significant alignment in data representation, highlighting a high degree of similarity in the orientation of the data vectors across the datasets.

Similarly, the Pearson correlation coefficients were notably high, ranging between 0.98 and 0.99. These strong positive correlations point to a robust and consistent linear relationship between each dataset pair, further confirming the parallelism in their data patterns.

In terms of the iLISI scores and their corresponding capacities, there was observed variation in integration quality across the dataset comparisons. The iLISI scores varied from 1.106 to 1.933, and capacities ranged from 1.356 to 1.984. These results highlight differing levels of integration efficacy, with the pairs scGenerated & scPBMC1 and scPBMC1 & scPBMC2 showing notably higher integration quality.

Lastly, the Correlation Discrepancy (CD) values, which ranged from 2.34 to 37.72, provided insights into the variability of gene expression correlation among the datasets. The lowest discrepancy was observed in the scGenerated & scPBMC1 comparison, suggesting a particularly close match in gene expression profiles. This result aligns with expectations, considering that PBMC1 and PBMC2 are experimental replicates of the bulk RNA-seq PBMC datasets and were trained on the same bulk2sc model, which was also trained on the paired scRNA-seq dataset.

Figure 10 in Supplementary Materials shows the QQ-plots for the distributions of the mean count data for the scRNA-seq data and the three generated single cell data generated from pseudo-bulk and real bulk datasets. The significant similarity in the demonstrated distributional patterns between the average generated count data to the average single cell count data suggest that the **bulk2sc** managed to learn accurate count data as a function of bulk RNA-seq data.

**Fig. 10.**
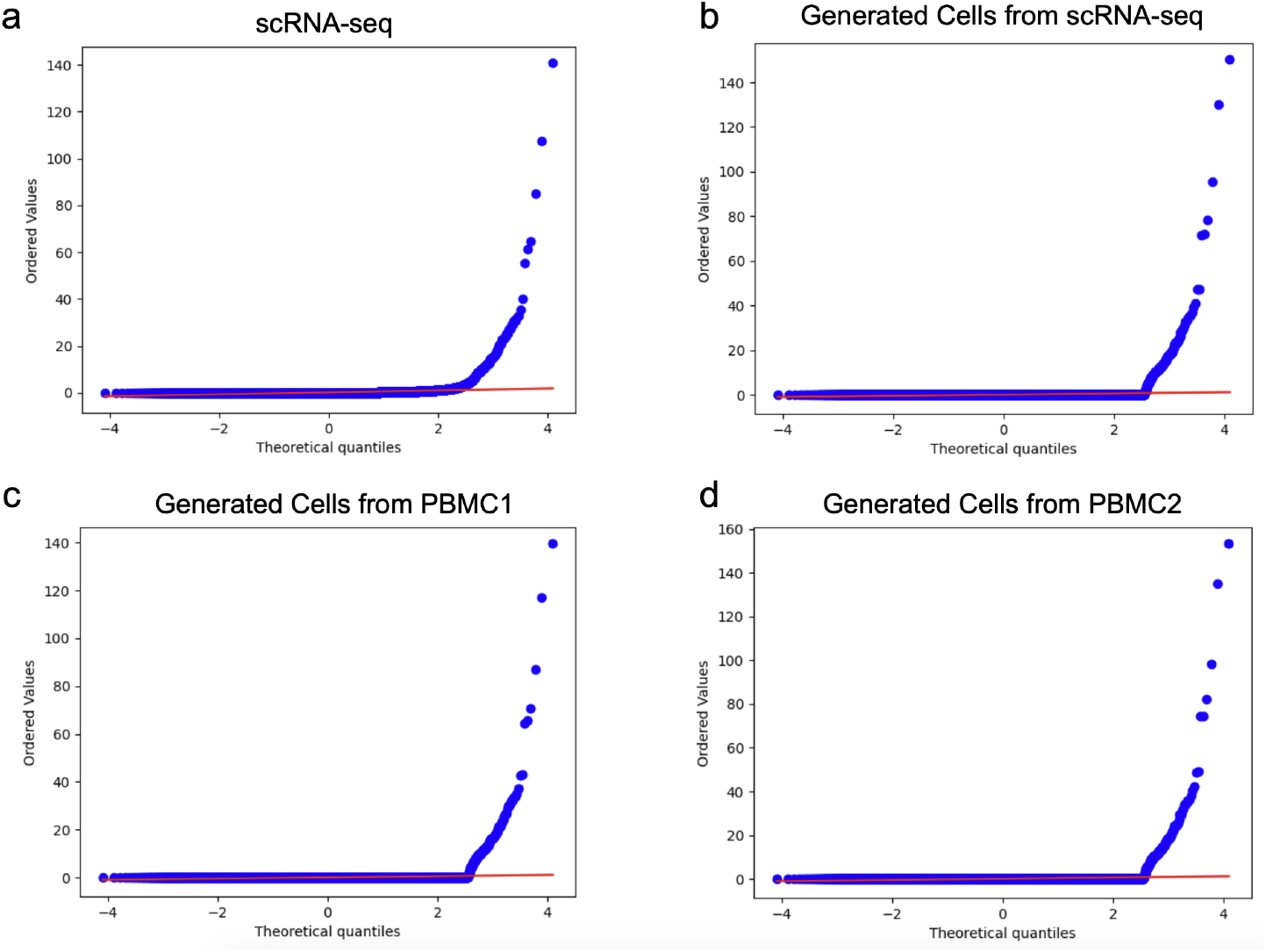
Quantile-Quantile (QQ) plots demonstrating the distributional characteristics of a. paired scRNA-seq data, b. generated cells from scRNA-seq data, c. paired bulk RNA-seq PBMC1 and d. paired bulk RNA-seq PBMC1.

### Data and Source Code Availability

All three PBMC datasets used in this study are publicly accessible on the 10X Genomics website. The paired scRNA-seq dataset and PBMC1 and PBMC2 bulk RNA-seq datasets can be found in (8). The source code for our model is available in the following Github repository: https://github.com/berkuva/Bulk2SingleCell.

## Conclusion

The high expense of single-cell data results in restricted sample sizes. Conversely, there is a vast amount of available bulk RNA-seq data. Our research introduces a new technique to derive single cell data from bulk RNA-seq datasets. Training Gaussian mixture parameters in a VAE’s latent space is known to be a difficult problem, so our method builds on previous research on parameter learning from reparameterization in this latent dimension of a VAE that is trained for dropout imputation. **bulk2sc** is trained to learn cell type-specific distributional parameters and proportions from single-cell RNA-seq data which are then learned by an encoder using bulk data as input. Our results indicate that **bulk2sc** can successfully generate representative scRNA-seq from bulk RNA-seq data with moderate to high degrees of similarity to the real data. Our overarching goal is to generate realistic single cell data for various tissues using bulk RNA-seq data. We are confident that our **bulk2sc** framework provides a solid foundation for generating single-cell data from bulk RNA-seq datasets and advances the role of generative machine learning in single cell research.

## Supplementary Materials

In Figure 2 in the main manuscript and Figures 4 and 5 below, we show the UMAP (Uniform Manifold Approximation and Projection) visualizations of five different stages of **bulk2sc**. Namely, they are the input scRNA-seq data (panel a), the reparameterized latent representation *z* (panel b), the reconstructed data from scGMVAE (panel c), the generated cells from the pseudo-bulk data constructed using the training dataset (80% of input cells) (panel d), and the generated cells from the pseudo-bulk data constructed using the testing dataset (20% of input cells) (panel e). In the five panels, distinct clusters corresponding to specific cell types are evident, closely mirroring the cell type proportions found in the scRNA-seq data, with minimal overlap observed. The consistency in cell type proportions between the scRNA-seq input datasets and the generated datasets is notably well-preserved, as demonstrated in the Quantile-Quantile (QQ) plots detail in Figure 6. These two Figures together demonstrates **bulk2sc**’s ability to not only generate cells with the type-specific characteristics accurately but also to maintain the relative abundance of each cell type.

We utilize QQ plots as a method for qualitatively assessing the alignment of probability distributions across distinct datasets: the input data, the single-cell data derived from the training set, and the single-cell data derived from the test set. These QQ plots offer a detailed view of the distributional comparison between the model-generated single-cell data and the original scRNA-seq data. This comparison is critical for evaluating the extent to which the model accurately replicates the statistical characteristics inherent to the sparse count nature of scRNA-seq data. Such an analysis is crucial for confirming the model’s effectiveness in preserving distributional consistency throughout both the training and testing phases.

In Figure 7, we present the QQ plots that demonstrates how well **bulk2sc** is able to model the distributional characteristics of the input scRNA-seq data in the generated single cell datasets from pseudo-bulk datasets formed by the training and the testing datasets. Specifically, panels a, d, g demonstrate the average gene counts in input scRNA-seq datasets with respective sample sizes of 3K, 10K, and 68K PBMC. Panels b, e, h exhibit the average gene counts for scRNA-seq data generated from pseudo-bulk RNA-seq data using the training set, while panels c, f, i depict those generated using the test set. It is clear from the QQ plots that the average gene expression values are highly similar between the scRNA-seq input datasets and the generated single cell datasets.

In the study conducted in (8), single-cell RNA sequencing (scRNA-seq) was used to explore and compare various sequencing methodologies (i.e. Smart-seq2 (30), CEL-Seq2 (11), 10x Chromium (versions v2 and v3) (36), Drop-seq (24), Seq-Well (9), inDrops (17), and sci-RNA-seq (5)) using different sample types, including human peripheral blood mononuclear cells (PBMCs). PBMC1 and PBMC2 represented two separate experimental replicates using PBMCs. These replicate experiments were conducted for assessing the reproducibility and reliability of the scRNA-seq methods under examination.

We generated single cell data from bulk RNA-seq datasets and compared the result with the paired scRNA-seq data as well as with the generated cells from the scRNA-seq data. Then, we applied the identical set of evaluation metrics to assess the similarities between the scRNA-seq data, the data generated from psedu-bulk RNA-seq data, and the data generated from PBMC1 and PBMC2 bulk RNA-seq datasets.

In Figure 8, we provide UMAP visualizations of the scRNA-seq data (panel a), various stages in training our **bulk2sc** model, such as the reparameterized latent representation *z* in the scGMVAE model (panel b), the reconstructed data by the scGMVAE model (panel c), and the generated cell from genVAE (panel d). Panels e and f represent UMAPs of single cell data generated from bulk RNA-seq data, PBMC1 and PBMC2 respectively. Numeric labels indicate Leiden (31) cell clusters, which are also highlighted in unique colors. N/A represents cell types generated that were not present in the single cell RNA-seq data.

In the presented UMAP visualizations in Figure 8, distinct clusters corresponding to Leiden-defined cell types are evident within the reparameterized latent space representation, denoted as *z*. These clusters become more defined in the reconstructed data, and this refined structure is preserved in the data generated from scRNA-seq. Notably, single cells derived from the actual bulk datasets, specifically PBMC1 and PBMC2, display a high degree of similarity to the cell type-specific clusters observed in the UMAP of cells generated from scRNA-seq.

Figure 9 compare the cell type proportions in the scRNA-seq data (blue), the generated cells from scRNA-seq data (orange), the generated cells from PBMC1 bulk RNA-seq data (green), and the generated cells from PBMC1 bulk RNA-seq data (pink). As shown clearly, the proportions in all synthetic single cell data follow the proportions in the scRNA-seq data, which indicates **bulk2sc**’s ability to model cell type information well from bulk RNA-seq data.

Figure 10 provides a comparative analysis of distributional characteristics across the four aforementioned four data sets in QQ-plots. Panel (a) displays the distribution of average count data in the original scRNA-seq data. Panel (b) shows the distribution for cells generated from scRNA-seq data using the **bulk2sc** method. In panel (c), illustrates the distribution of single-cell data generated by applying **bulk2sc** to PBMC1 bulk RNA-seq data. Finally, panel (d) illustrates the distribution of single-cell data derived from PBMC2 bulk RNA-seq data using the **bulk2sc** method. A notable observation across all four QQ-plots is the remarkable similarity in distributional patterns, indicating that the average count data across these different methods and data sets remains consistently close to each other, which highlights **bulk2sc**’s ability to closely model the count data from both pseudo-bulk and true bulk RNA-seq datasets.

We present the quantitative analyses in Table 5. In our analysis, cosine similarity scores consistently showed high values from 0.97 to 0.99 across datasets, indicating strong alignment in data representation. Pearson correlation coefficients echoed this trend with impressive values from 0.98 to 0.99, confirming a robust linear relationship between dataset pairs.

The Integration Local Inverse Simpson’s Index (iLISI) scores varied, with ranges from 1.106 to 1.933, and capacities from 1.356 to 1.984, highlighting differences in integration effectiveness. Specifically, the scGenerated & scPBMC1 and scPBMC1 & scPBMC2 pairs demonstrated superior integration quality.

Furthermore, the Correlation Discrepancy (CD) values, which are calculated as the 1-norm of the difference in correlations to gauge the discrepancy between them, varied from 2.34 to 37.72. This range underscores the degree of variability in gene expression correlation across the datasets. The smallest discrepancy in the scGenerated & scPBMC1 pair suggested a close match in gene expression, consistent with the use of the same bulk2sc model for PBMC1 and PBMC2, which are replicates of the bulk RNA-seq PBMC datasets.

